# Science in School: Transforming K-12 Outreach through Scientist Teacher Partnerships

**DOI:** 10.1101/2021.07.27.453770

**Authors:** Brian Abramowitz, Megan Ennes, Stephanie Killingsworth, Pavlo Antonenko, Bruce MacFadden, Alan Ivory

## Abstract

The Scientist in Every Florida School (SEFS) program was started in 2019 with a long-term vision to connect Earth systems scientists with public K-12 schools in Florida and therefore create long-term scientist-teacher partnerships. SEFS fulfills personalized requests to create meaningful and impactful interactions to support teacher pedagogy and student learning. We have as part of our mission a focus on mainstream public schools, and in particular, those that are Title I. We also are committed to working with at-risk teachers. The major components of our program include the scientist-teacher partnerships, focused professional development workshops on Florida’s Earth systems (air, water, land, and life), classroom visits, and other web-based activities. Although still only in its first few years, the project and its more than 600 scientists have a wide reach with over 850 teachers and 53,000 students participating in our programs, which were delivered virtually in the 2020-2021 school year covering about 60% of Florida’s 67 counties. In this article, we describe our programmatic features as well as recommendations for those who could implement similar programs.

---

Florida is on the front lines of massive Earth systems changes that are threatening the environment, economy, and our way of life. Earth systems refer to the interaction of air (atmosphere), water (hydrosphere), land (geosphere), and life (biosphere), and how humans impact them. Current research indicates that educating the next generation about societally relevant issues such as climate change is vital for developing a scientifically literate public empowered to take action to address these issues (e.g., Hahn, 2021; Peterson, et al., 2019). This emphasis will hopefully lead to greater change because K-12 learners are likely more receptive than adults to learning about these topics (Stevenson et al., 2014; Kahan, 2012). Unfortunately, science often takes the backseat in the K-12 system, despite its contribution to social and economic progress (Marincola, 2006). This is particularly true in elementary schools where some teachers lack the background, confidence, or mandate to teach science (Menon & Sadler, 2016). Science in middle and high schools can also suffer from variable and outdated content knowledge and inconsistent teacher pedagogy (Park & Oliver, 2008).

To help support high-quality science teaching through scientist-teacher partnerships, the Scientist in Every Florida School (SEFS) program was developed in the fall of 2018. The University of Florida’s (UF) Thompson Earth Systems Institute received funding from the UF President’s office along with support from other stakeholders to fund this as a pilot program for four years. This set the foundation for their “Moonshot” initiative entitled “Scientist in Every Florida School’’ (Spence, 2019). The concept of a “moonshot” comes from President John F. Kennedy, who in the early 1960s presented a lofty goal--to put an astronaut on the moon by the end of that decade (Brinkley, 2019; Oliphant 2019). This was a seemingly unattainable challenge because the science and technology infrastructure of the time was not prepared to do this. Thus, during the remainder of the 1960s, many aspects of this goal were achieved through scientific and technological innovations. In the same vein, our SEFS program seeks to provide innovation and develop new ways and best practices for reaching Florida schools, their teachers, and students. The mission of SEFS is to engage Florida’s K-12 students and teachers in cutting-edge science in a way that provides role models and experiences to inspire the future stewards of our planet. Scientist role models include graduate students, post-doctoral researchers, university faculty, and industry partners at local, state, and federal agencies. Program participation helps teachers feel more confident teaching content after talking to an expert. Additionally, students feel inspired by the role models who visit their class, will be more aware of Florida’s environmental changes around them resulting from climate change, and will learn what can be done about these issues. In this article, we describe our program and what we have learned so far as we move forward with our decadal vision of impacting Florida science education in public schools.

SEFS goals are to build long-term collaborative relationships between teachers and scientists, introduce K-12 students to scientists in a wide range of Earth systems careers, and increase the integration of current scientific research and big data into classroom lessons. To achieve these goals, SEFS coordinates virtual and in-person scientist visits to schools across Florida. These visits serve to engage students with real-world Earth systems science and increase interest in related science, technology, engineering, and math (STEM) careers. Although other similar programs exist, e.g., Bay Area Scientists Inspiring Students (BASIS, n.d.), Skype a Scientist (2021), and Letter to a Pre-Scientist (n.d.), SEFS is distinguished by its unique statewide initiative that prioritizes long-term relationships through personalized interactions via scientist-teacher partnerships, customized school visits, and an emphasis on public schools. We place priority on schools that are designated as Title I, which is a federal program that provides financial assistance to schools for children from low-income families to help ensure that they meet challenging state academic standards (NCES, 2021).

## Vision and rationale

As the name connotes, over the next decade our long-term goal is to have scientists “visit,” “reach,” or “impact,” every K-12 public school in Florida at least once a year. With more than 3,000 public schools in the 67 Florida counties (School, n.d.), this is indeed a daunting task, but it remains our aspirational vision--one by virtue of its title that is easily communicated and quickly grasped by stakeholders.

In the preceding paragraph, we use three terms: visit, reach, and impact, each of which has a specific meaning. We frequently use these as activity metrics in the context of the SEFS program. A *visit* is a discrete event in which a scientist goes into a classroom, either in-person or virtually. We typically gather data on the number of teachers and students, as well as which scientists participated at a particular visit. *Reach* is used to quantify the number of teachers or students to whom we presented our content. We might say, for example, that during a particular professional development (PD) session we interacted with 70 teachers, meaning that we counted the attendees as having been reached through our activities. In terms of efficacy, however, neither visit nor reach allows us to understand what our audiences learned or how they benefited from our SEFS activities. In contrast, *impact* is a term reserved to describe how we improved engagement, understanding, and/or learning. We assess impact via evaluation, primarily through teacher surveys, but we also pay attention to qualitative feedback from participants, e.g., open-ended survey questions or unsolicited email responses. In our program, impact is the best indicator of the efficacy of our programs and activities; on the other hand, it is also the most time-consuming of our activities to gather relevant data.

Our program activities ramped up significantly in July of 2019 with the hiring of two K-12 SEFS coordinators (authors Abramowitz and Killingsworth) and our first annual summer PD. Between July 2019 and February 2020 we developed a comprehensive slate of in-person classroom visits, district-focused PDs, and web-based activities. During this phase, we felt that our in-person classroom activities were a hallmark of our program. However, in early 2020, with the onset of COVID-19, physical schools closed down and we quickly pivoted to fully virtual programming for all of our activities. Given what we have learned during COVID-19, our transformation to mostly virtual programming allowed us to both scale up our programming and economize our efforts, for example, the cost of travel was greatly reduced.

## Strategic Components and Goals

Through formal strategic planning, we have identified several related goals for the SEFS program that, taken together with the overall vision, comprise the unique scope of our project.

### 1. Program focus

SEFS is a program within the Thompson Earth Systems Institute (TESI, 2021) at the UF Florida Museum of Natural History. The mission of TESI is to enhance understanding of Earth systems including the atmosphere, hydrosphere, geosphere, and biosphere as well as the impact of humans on these systems. Simply put, we use the phrase “air, water, land, and life” as our thematic scope. When making decisions about programming and resource allocation, we are careful to focus only on these content domains. For example, we do not specifically focus on health-related content. Another theme of our programmatic activities includes the Nature of Science because this is something that teachers are expected to teach and many Florida Standards align with this concept (Welcome to CPALMS, 2019). Repeated feedback from teachers indicates the need for students to better understand the process of science as it applies to different content domains.

### 2. Target audience

SEFS is envisioned to serve mainstream public schools in Florida. Our rationale stems from the mission of UF as a land-grant, publicly funded university, and our general commitment to public education. With regard to specific activities and programs (e.g., school visits), we therefore tend not to focus on charter schools, private schools, and home-schooling, or the recent rise of K-12 learning pods (e.g., Sergent, 2020). However, when we have space available in our programs such as our teacher PD opportunities, we make them available to these groups. Likewise, some of our webinars have broader intended audiences. For example, when we partner with informal learning institutions such as museums, nature centers, and botanical gardens, our virtual audience might include a broader K-12 composition as well as lifelong learners.

### 3. Scientist-teacher partnerships

Our teacher-focused activities include classroom support via scientist visits and discrete PD (Figure 1). Rather than a one-off for either of these activities, SEFS aspires to build lasting relationships between the scientists and teachers. We envision that teachers will view the scientist as an available content resource provider and can call upon them as makes sense. We believe that this strategy not only is important to the teachers, but as other studies have shown, these scientist-teacher partnerships also benefit the scientists (e.g., Tanner, 2000; MacFadden 2019).

**Figure 1.** Scientists are mentoring teachers who take on the role of a scientist as they are provided an authentic lab experience

### 4. *Diversity, Equity, Accessibility, and Inclusion* (DEAI)

Part of our program plan is to develop activities and reach schools that promote DEAI. In addition to this being part of our TESI core culture, we do this in two specific ways: (a) within the K-12 districts that we have served so far, we place priority on Title I schools and their teachers, and (b) we deliberately select scientists who exemplify diverse and near-peer role models. Related to the latter, some teachers request that the scientists coming into their school share similar backgrounds with the students.

### 5. At-risk teachers

Retention of public-school teachers is widely recognized as a systemic challenge affecting the K-12 educational workforce throughout the U.S., and so too is the case in Florida (García & Weiss, 2016). An example of this is in January 2019, with half of the K-12 school year completed, Florida schools reported over 2,200 instructional positions still unfilled. Further, the vacancy with the greatest need was science. While schools were able to actively recruit to partially fill this gap, some schools had to hire substitute teachers (Isger, 2019). The effects of this attrition are profound and relate to overall instructional stability and student achievement. We define at-risk teachers as those new to teaching (those in their first five years), or those in their mid-career and considering leaving the profession (also see Sparks, 2018).

### 6. Scientist in Residence

As a result of philanthropic support dedicated to rural counties in north-central Florida, SEFS has developed a partnership with the Marion County School District. The Marion County School District is mid-size with 48 schools, of which 44 are Title I. This opportunity allowed us to develop an innovative full-time scientist-in-residence program located at the Silver River Museum (2021), which is part of this public school district. This program, and COVID-19, allowed us to experiment with different content delivery systems. During the time that the schools were either fully or partially closed to in-person instruction, our scientist-in-residence (author Ivory) conducted web-based virtual STEM learning to specific classrooms as well as combined presentations. The SEFS scientist in residence program resulted in a far greater reach in Marion County relative to other counties, while the virtual programming resulted in a better ability for us to scale up our reach (Figure 2). Now that we are transitioning to more in-person activities, we also are using the newly implemented science lab at the Silver River Museum for our programming which will include more active learning and hands-on activities, e.g., related to local water quality.

**Figure 2.**
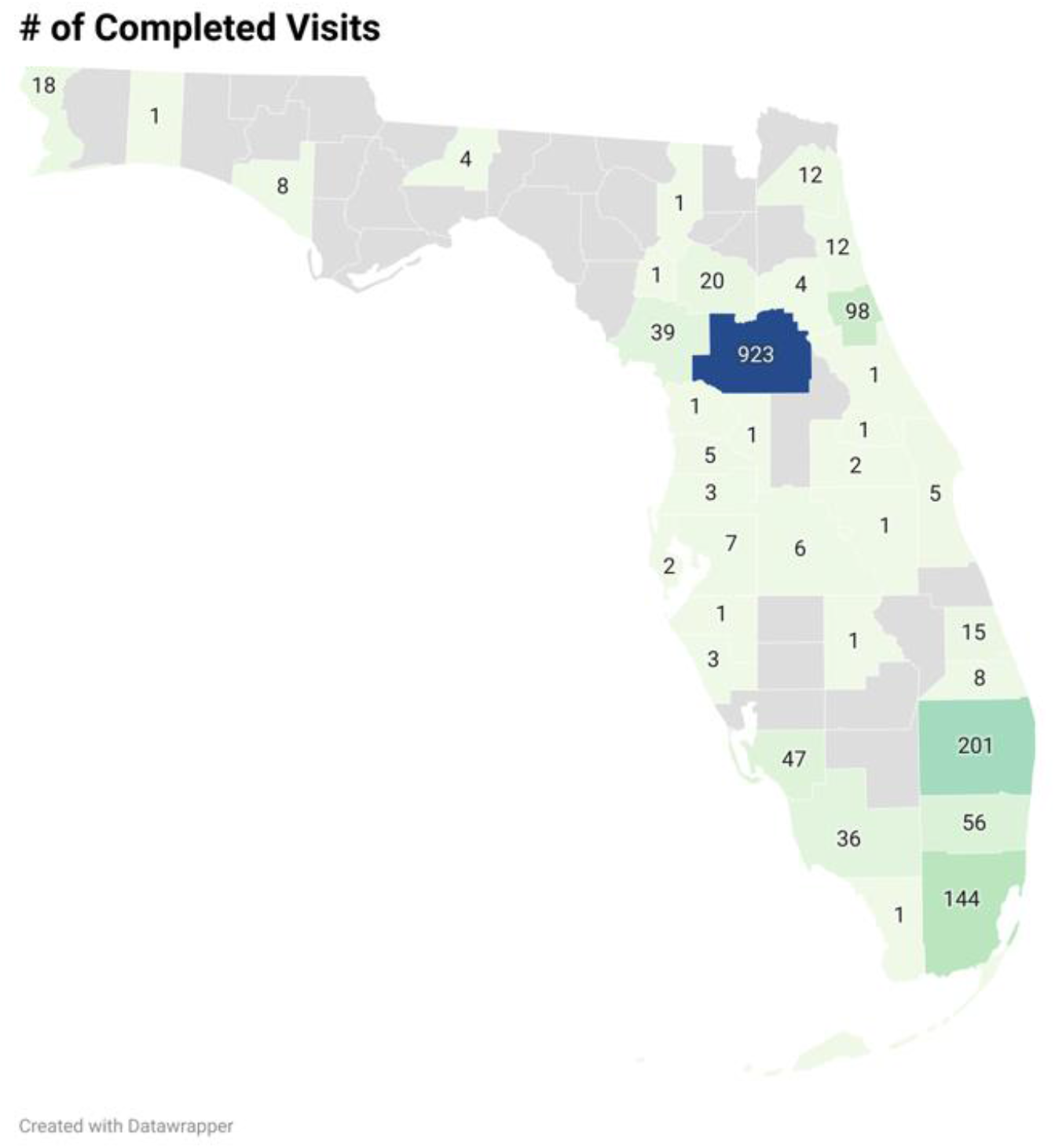
The Number of Completed Scientist visits in Florida during the 2020-2021 school year.

## Conceptual Framework

Today, K-12 science education in the U.S. is informed by the Framework for K-12 Science Education released by the US National Research Council in 2012 (National Research Council, 2012). This framework is often referred to by K-12 educators and researchers as the 3D science learning framework because it conceptualizes K-12 science education as consisting of three core dimensions: a) STEM practices, b) crosscutting concepts, and c) disciplinary core ideas. Furthermore, while science teachers have numerous resources such as curricula and textbooks for incorporating crosscutting concepts (e.g., structure and function) and disciplinary core ideas (e.g., photosynthesis) in their instruction, integration of authentic scientific practices in science instruction presents more of a challenge. The main reason for this is that until recently teacher preparation programs, curricula, and methods have focused primarily on the dimensions of disciplinary core ideas and crosscutting concepts, whereas the practices of doing science and STEM more generally were left largely unaddressed (National Research Council, 2012, 2014).

The SEFS program addresses the issue of improving science teachers’ understanding and integration of STEM practices in their instruction by matching them with scientists who can demonstrate in K-12 classrooms not only the science they do and why but also *how* they go about doing it: making clear the tacit STEM practices scientists use to address important societal issues. This focus on science as a set of core STEM practices is particularly important now when many of these fields rely heavily on technology, mathematics, and engineering. For example, paleontology organically integrates concepts and content from biology, environmental science, geology, oceanography, anthropology, while also harnessing the resources and tools available from other fields of STEM, including technology (e.g., big data in the cloud, 3D scanning; Callaway, 2011), engineering (e.g., advanced analytical 3D imaging, digital manufacturing; Hooper, 2013), and complex mathematical modeling, statistical algorithms, and machine learning (Elewa, 2011).

As a PD program at its core, SEFS uses best practices of teacher PD. Recent research found that effective PD has seven characteristics (Darling-Hammond et al., 2017). These include a focus on content, active learning, support for collaboration, models of effective practice, coaching and expert support, feedback/reflection, and a sustained duration (Bates & Morgan, 2018). Our program includes each of these components as described below (Figure 3).

**Figure 3.**
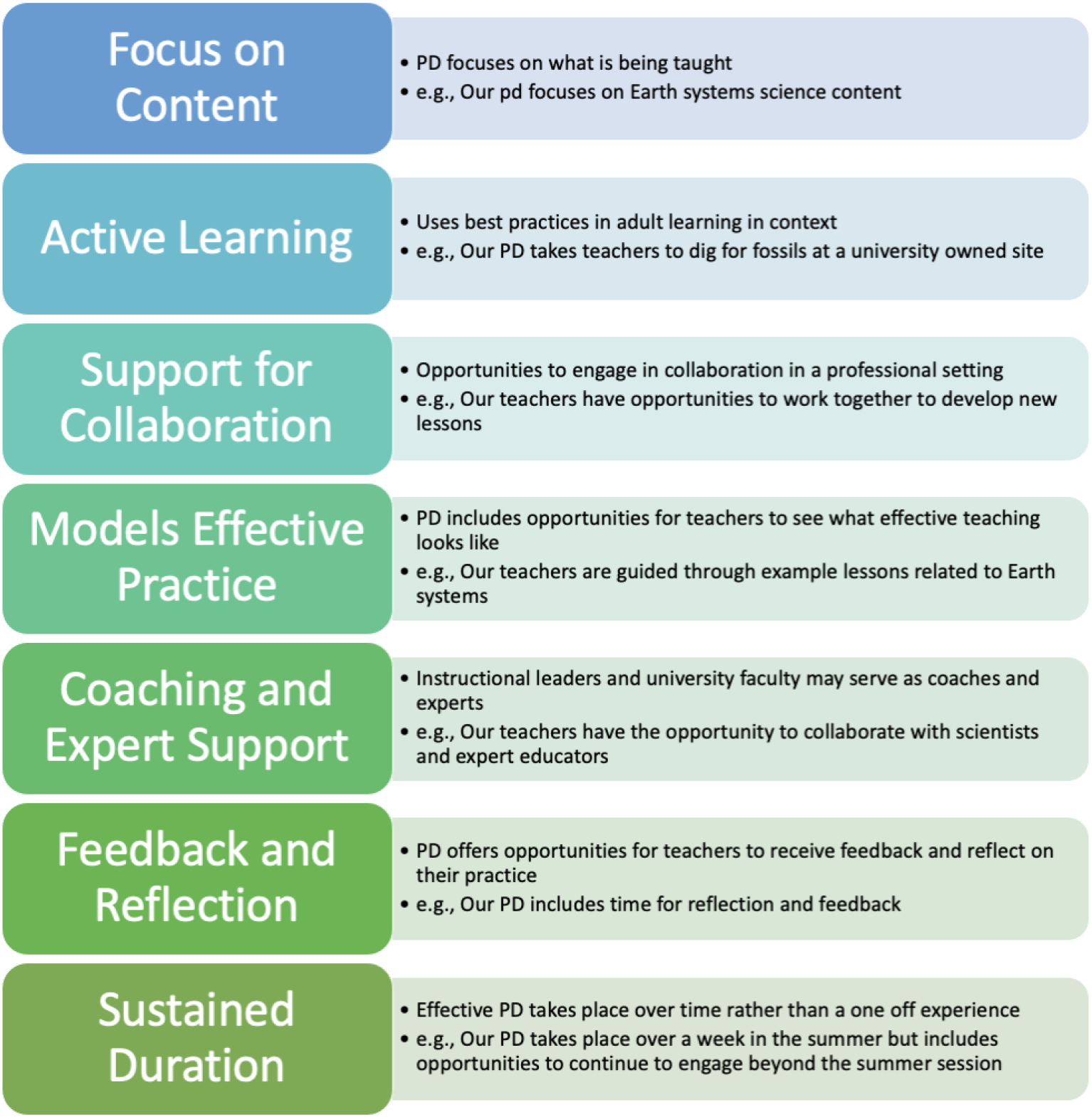
Characteristics of effective professional development for teachers as described by Bates & Morgan, 2018.

### Focus on content

Each year, the PD includes an area of content focus that relates to Earth systems science such as the biosphere (2019), nature of science (2020), and hydrosphere (2021). Content delivery includes keynotes and short presentations (“sparks”) by experts in the respective fields, deeper dives into content with discussion, breakout sessions, and Q&A with content experts, along with hands-on research experiences, field trips, and lesson collaboration. The K-12 education and outreach coordinators work with the scientists to better prepare them for connecting with and teaching content to teachers. This involves identifying and aligning relevant state standards related to their area of research.

### Active learning

To involve our participants in active learning, the PD model “engages educators using authentic artifacts, interactive activities, and other strategies to provide deeply embedded, highly contextualized professional learning” (Darling-Hammond et al., 2017, p. 7). In non-COVID-19 years, PD includes authentic research experiences in the lab and field for teachers, field trips to nearby localities with expert guides, materials, and supplies for lessons to be implemented in the classroom, as well as ample time for small group and whole group discussion, collaboration, and reflection. For example, a group of teachers was taken to a UF fossil site where a fifth-grade teacher from Pensacola, Florida, discovered a five-million-year-old fossil horse tooth of scientific value (Burton, 2019). The newly uncovered fossil was added to the collections at the Florida Museum of Natural History, and as such, the teachers participated in authentic research discoveries. Since the onset of COVID-19 in March 2020, SEFS PDs have remained virtual and teachers and scientists connect remotely. If needed, supplies are shipped to the teacher for hands-on activities beforehand so they remain an integral part of the experience. The SEFS team iteratively reviews the workshop program and agenda and we make modifications based on survey feedback and debriefing notes.

### Support for collaboration

Collaboration allows teachers to break free of the structured environment of the classroom as well as to encourage change beyond an individual classroom (Darling-Hammond et al., 2017). To encourage collaboration, teachers work in groups with the support of their partner scientist throughout the week to develop lessons and extension activities to be taught in the classroom during the following school year. As part of the process, teachers are responsible for scheduling a scientist visit for their classrooms the following year and sharing artifacts with the SEFS team from the lesson implementation.

Collaboration elements are infused in the PD in several ways to make workshops feel like a safe and inclusive environment for teachers to feel comfortable as learners, to explore, and ask questions. Teachers participate in a meet and greet before the workshop and regularly connect as a whole group as well as in smaller breakout groups during the week. They are also invited to participate in the SEFS Facebook group (a professional learning community and network of Florida teachers and scientists). Additionally, we encourage them to reach out to the SEFS team and their partner scientist at any time. Many teachers are invited back for subsequent PD opportunities and later teacher cohorts as teacher leaders. All of these collaboration opportunities allow for sustained social learning.

### Models of effective practice

Modeling of activities, concepts, and the process of science are fundamental to our PD workshops. These practices help teachers to visualize how they can implement new strategies in their own classroom (Darling-Hammond et al., 2017). The SEFS team models a related activity with the teachers who then take time to explore the activity themselves or in groups. Additionally, teachers come to the PD with their learning standards (Welcome to CPALMS, 2019) and pacing guides to best integrate the content learned. Collaboration with other teachers from the same grade bands and with scientists helps facilitate the creation of novel and innovative lessons and activities that incorporate the current research and content being taught with the standards taught in K-12 classrooms. The SEFS team often guides or supports teachers in this process. Teachers get to experience science first-hand which makes them not only more comfortable with the content but more confident and excited to teach it to their students.

### Coaching and expert support

Experts have an important role in guiding and facilitating teachers’ learning as it relates to their classroom practice (Darling-Hammond et al., 2017). Our program model relies on expert educators as facilitators and role model science experts to deliver current scientific research and understanding in a way that is accurate and trustworthy. Both teachers and students benefit from these role model interactions to create rich learning opportunities and inspire the next generation of Florida’s environmental stewards. District leaders also often participate in the SEFS PDs so that they can provide additional support and feedback to their teachers who attended.

### Feedback and reflection

Effective PD allows teachers opportunities to engage in reflection, receive input related to their practice, and make changes as needed (Darling-Hammond et al., 2017). To address this need, the PD agenda has reflective time built in each day. This helps the participants gain valuable feedback and for program organizers to gauge the teachers’ level of understanding and aids in pacing as well as the depth and breadth of content delivery at the PD.

### Sustained duration

Best practices in teacher PD underline the importance of sustained engagement between the teachers that offers teachers multiple opportunities to learn about the content (Darling-Hammond et al., 2017). In addition to the week-long PD experience, teachers who participate frequently remain engaged in additional PD opportunities. They typically apply to attend subsequent workshops, make multiple scientist requests for virtual and physical classroom visits throughout the year, and frequently attend the SEFS livestream programming. Teacher surveys have provided feedback to the team that teacher participation has sharpened their understanding of content, provided new ideas for lessons, and perhaps most importantly, has re-energized them to teach.

## Evaluation & Data Collection

Evaluation of the efficacy of the program to engage program participants in the teacher-scientist partnership is accomplished using the custom-designed Scientist-Teacher Educational Partnership Survey (STEPS, Appendix A). STEPS consists of 15 items that are completed independently by both the teacher and the scientist. The three factors - Communication and Planning, Science Teaching Self-Efficacy, and Perceptions of Student Engagement and Learning - help the scientist and the teacher partners independently reflect on the important aspects of the partnership experience. Once SEFS collects an adequate number of teacher and scientist STEPS survey data, we will perform a confirmatory factor analysis to determine whether the items load onto the three factors that underlie the survey design. At this stage of the program implementation, at least 75% of the teacher and scientist respondents either agree or strongly agree that a) the communication between the partners was timely and helpful and b) they are learning new and useful strategies for planning and implementing science instruction. More than 90% of the respondents believe that students were engaged and interested during the SEFS-sponsored educational events and enjoyed the interaction with the scientist.

## Benefits to Teachers/Scientists

Evidence indicates that participation in scientist-teacher partnerships can have positive effects on the scientists, teachers, and students (e.g., Brown et al., 2014; Herrington, et al., 2016; Shein & Tsai, 2015; Ufnar et al., 2017, 2018). One study found that participation in such a program resulted in positive belief, value, and attitude changes for teachers regarding their ability to teach inquiry-based science in their classrooms and their schools (Herrington et al., 2016). Another study found that scientist-teacher partnerships not only increased the teachers’ ability to teach via inquiry but also increased their confidence and content knowledge (Ufnar et al., 2017). This has also been found globally as a study in Taiwan found that participation increased teachers’ levels of content knowledge as well as their pedagogical content knowledge (PCK) and the scientists felt they had a better understanding of how students learn, what is taught in schools, and PCK (Shein & Tsai, 2015).

While many studies focus on the impacts of scientist-teacher partnerships on students and teachers, there are a few that focus on scientists (e.g. Tanner, 2000; Ufnar et al., 2017). A study of 34 scientists found that the program benefited scientists as professionals, university educators, and individuals (Tanner, 2000). Benefits to scientists as professionals included the ability to engage with their colleagues in new ways, the development of new skills that are useful in their line of work, the ability to reflect on their understanding of or excitement related to science, and the opportunity to consider new careers (Tanner, 2000). As future educators, the benefits included learning to explain content more simply, learning about pedagogy related to teaching science, and the ability to put these strategies into practice. They also learned more about teachers and students in K-12 schools and concluded that they can act as student role models. Finally, through this participation, individuals increased their self-efficacy, were able to engage in community service, and they saw an increase in satisfaction that enriched their lives (Tanner, 2000). Similarly, the study by Ufnar and colleagues (2017) found that participation allowed scientists to gain skills related to communication, mentoring, and teaching as well as allowing them to serve as student role models.

## Logistics

The program has two K-12 education and outreach coordinators who streamline and fully support the process of a scientist’s classroom visit. By having outreach coordinators employed by the program, the burden and/or commitment of time, as well as K-12 pedagogical teaching expertise required by scientists, is greatly reduced (Andrews et al., 2005). The process of creating partnerships between teachers and scientists and preparing scientists for the classroom interaction is one of the primary roles of the coordinators.

The scientist classroom visit request protocol follows a specific process. The coordinators have created a Google form that prompts the teacher for information about their vision for the visit. These details assist in the coordinator’s identification of a scientist who will be the best match for the request. This form can be found on the SEFS website, Facebook group, monthly newsletter, Florida school district websites and promotional messages, and on the Florida Association of Science Teachers website (Florida Association of Science Teachers, n.d.). Once submitted, the coordinator who oversees that Florida county will confirm receipt of the request with the teacher and then begin reaching out to potential scientist participants. When a match is found, the coordinator will introduce the scientist and teacher to create a long-term scientist-teacher partnership. Included in this introduction email, both teacher and scientist receive an infographic with research-based best practices for a successful visit. The scientist receives a SEFS branded PowerPoint slide deck for use during the classroom interaction. Also, both parties involved will be encouraged to further discuss the details of the planned visit (including student prior knowledge, defining the learning standard, or other logistics). The email emphasizes keeping a coordinator copied on the emails so they can be kept aware of what is going on and support the visit at each step of the process. Additionally, if travel accommodations, such as hotel and car rental are required, then a coordinator will arrange and pay for this aspect of the visit.

After the completion of either the physical or virtual classroom visit, an electronic survey is distributed to the teacher to assess the quality of the program. The results indicate areas of success and possible areas to check in on with the scientists for continued science communication growth. The information provided also informs the coordinators if any changes need to be made during the scientist preparation and training phase. To continue building the program, coordinators may reach out to the teacher who completed the survey to use their text as a testimonial.

## Barriers, Challenges, and Lessons Learned

The global COVID-19 pandemic has been a major factor affecting our SEFS programming and outreach. Thus in March 2020, all in-person classroom visits and university travel were closed down and largely remained that way through July 2021. The SEFS team, therefore, quickly adapted by changing the presentation format of all of the programming to virtual delivery, whether it be multi-class presentations, single classroom virtual visits, or teacher PD.

We have learned many things from COVID-19, for example: (1) SEFS can remain active and in high demand regardless of whether the programming is virtual or in-person and (2) the program’s reach extends to many more classrooms virtually (Figure 4). The virtual delivery is done primarily via Zoom, but accommodations can be made for other platforms, such as Microsoft Teams and Google Meet, depending upon the needs of the particular school district.

**Figure 4.**
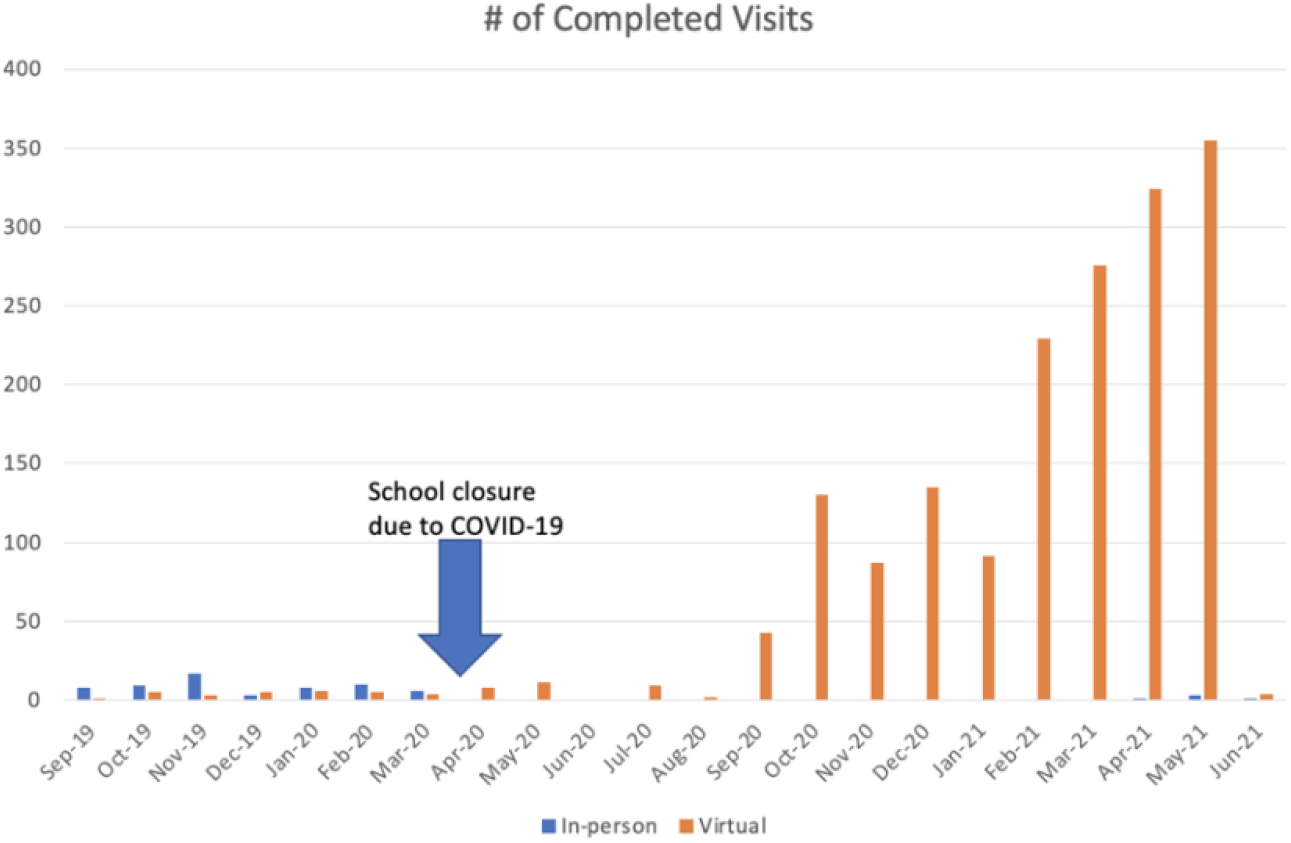
The number of completed visits from the onset of the SEFS program to the present.

Further, SEFS staff needed to provide additional technical support to Florida teachers. The sudden switch to virtual and/or hybrid teaching shifted pedagogical and classroom management demands. Both districts and teachers quickly realized that this huge alteration to teaching could be a barrier to student learning. Additionally, outside educational agencies and organizations that customarily hosted K-12 field trips were in effect grounded from outreach. Because of SEFS’s established virtual platform, some of these agencies looked to SEFS to partner and collaborate to deliver content. Thus, even during COVID-19, the demand for SEFS programming remained high, indeed it is ever-increasing throughout Florida.

While there is no substitute for the in-person classroom experience for the scientists, teachers, or students, the move to virtual has greatly increased the ability for SEFS to reach a greater number of K-12 audiences. From a strategic perspective, it has allowed the program to better understand how to be a resilient organization. This increased reach through the shift to virtual programming demonstrates the scalability of SEFS programs using virtual technology.

## Innovative Classroom Visit Examples

During the past two years, the SEFS program has witnessed many positive classroom visit experiences both in-person and virtual in classrooms around the state. Visits are at the request of teachers who share content goals for an interaction with a scientist. Often, visits aim to connect the standard-aligned content being taught in the classroom with real-world scientist role models and real-world applications. A frequent outcome of the teacher-scientist collaboration is an ongoing relationship with the classrooms. Several examples of innovative classroom visits throughout Florida are illustrated as follows:

- Often accessibility of lab equipment and supplies prevents teachers from offering authentic experiences for students that expose them to the tools and processes of science. Funding and lack of time can also be a deterrent for many teachers to provide field trip opportunities to their students as well (Tuthill & Klemm, 2002). As part of a genetics unit, teachers from a Title I high school in Palm Beach County wanted to expose their students to many of the common DNA techniques being used today to better understand inheritance. Students participated in a virtual field trip to the Equine Genetics Lab at UF. Their students watched the process of DNA extraction using horsehair and blood samples. A Palm Beach County teacher remarked,
  It’s truly fascinating to see how this process has changed in the last 15 years since I worked in a lab myself. I learned so much about how bioinformatics has changed the speed and volume with which DNA is sequenced, and it’s great to bring to light careers for my students.
  During the interaction, scientists from the UF Equine Genetics Lab also shared how artificial intelligence technology is being used to understand horse phenotypes for the horse racing industry. The visit gave students a chance to see the scientific process and common tools at use, the unpredictability of science, and all while touring a lab at UF, somewhere this Palm Beach group would be unable to visit normally due to the distance (Figure 5).

**Figure 5.** The UF Equine Genetics Lab demonstrates DNA extraction techniques for students across the state.
- As part of the scientist-in-residence program, in April 2021, through a virtual boat ride in the glass-bottom boat along the Silver River, our scientist reached 24 schools, 140 teachers, and 2,629 students. The glass-bottomed boats at the Silver Springs State Park are specifically designed to allow perfect viewing of the springs down below, but provide limited occupancy for patrons. This has made field trips on the glass-bottom boats a logistical and financial challenge for schools. In collaboration with the Silver Springs State Park, we were able to create a virtual glass-bottom boat tour for all K-5 Marion County students to enjoy. This was achieved by placing a camera on the glass itself, allowing the boat captain to share the rich history of the river, and letting the scientist explain what life is being shown through the glass. Exposing students to science in a tangible way shows them how interconnected science concepts can be to each other and the student themselves (Raved & Assaraf, 2010). For example, along the boat ride, students were introduced to the Florida aquifer and water cycle, the damage pollution has on their local waterways, invasive species, and even some of the research being conducted in that area. Rather than reading about these topics in a textbook, they saw how science plays out in the real world. A Marion County elementary school teacher said,
  Thank you for a great year of field trips! My students were very interested in what they could do to help and repeatedly asked if they could come clean up the river. They stayed on the edges of their seats and were sponges, absorbing everything they could.
- When a Duval County teacher was challenged to continue creating authentic science experiences for her students, she reached out to SEFS for support. Feeling nervous and unprepared to lead some activities on her own, she was hoping for a scientist to share their expertise. By establishing a teacher-scientist partnership, teachers can receive assistance which enables them to conquer their hesitations surrounding teaching certain content areas (Ufnar & Shepherd, 2018). After two weeks of collaborations and co-developing a lesson, the teacher and scientist began their classroom interaction. Although the visit was virtual due to COVID-19 restrictions, the 10 fourth-grade students were able to participate in an authentic hands-on experience just as they would with the scientist in the room. With the support of personal tablets, the scientists (in Tallahassee) and the students (in Jacksonville) were able to synchronously dissect squid specimens “together.” After reflecting on the experience, Florida State University’s Sea to See program scientist shared,
  Thank you both for giving us a chance to dissect squid with this wonderful class. [teacher] is obviously a fantastic teacher as evidenced by her student’s knowledge, engagement and the classroom culture of inquisitiveness. We had such a good time with this. Today I could see the smiles behind our masks everywhere I looked (Figure 6).

**Figure 6.** Sea-to-See Scientist virtually dissects squid with students.

## UF Courses and Related Instructional Activities

Being located at a public university, Ennes and MacFadden also serve as faculty in the Department of Natural History. Therefore, they have found ways to incorporate SEFS as components in their curriculum.

Ennes teaches an annual graduate course as part of the university’s Environmental Education and Communication certificate called Science Communication and Public Education. This course is designed to help future scientists translate their research for the public. One of the major assignments of this course was to film a video for the SEFS Science Segments. These short videos introduce K-12 students to diverse scientists who describe their research and how it is related to Florida’s state science standards. The students in her class also designed and practiced hands-on activities related to their research that can be implemented with programs such as SEFS. The student evaluations suggested that the class is meeting its goals of supporting scientist engagement with the public and K-12 audiences. Many were excited to use their new skills once it is safe to return to the classroom.

MacFadden teaches a graduate seminar once a year entitled Broader Impacts of Science on Society (also see MacFadden, 2019). A module within this course is devoted to K-12 outreach, with a focus on SEFS. Likewise, the course typically has guest talks, e.g., by our SEFS coordinators Abramowitz and Killingsworth about their prior experience as teachers and also their current work with SEFS. An expectation of this graduate seminar is a final class project, and over the years some of our students have elected to focus on K-12 outreach and SEFS.

## Funding and Business Model

The SEFS program was established through a four-year-long moonshot grant funded by the UF President’s Office in 2018. This was initially proposed to be a small pilot in three Florida counties that quickly grew to five, and after COVID-19 hit, the SEFS program expanded the number of counties that we have reached to almost three dozen by early 2021. Given this rapid expansion, the base level of support was quickly outgrown. Thus, to reach our current level of effort and build capacity, we have added to our funding stream. This mostly includes matching and other funds from other UF stakeholders (Provost, VP for Research, and Museum Director’s Office), income generated from a TESI endowment, and philanthropic gifts from private donors. The salaries of the three faculty in the TESI core team are contributed as a match from the Museum’s annual budget appropriations.

Our long-term business model is to offer our programs free of charge to the teachers and schools, with the staff salaries and operating expenses derived from the funding sources described above. We are also promoting SEFS as a program that can help scientists who submit proposals to federal grant agencies with a K-12 component of their Broader Impacts plan (MacFadden, 2019), and in so doing, include salary for our SEFS coordinators effort written into the scientists’ budgets. We also plan to provide program evaluation services as a cost-generating measure that would provide revenue back to sustain SEFS. Ultimately, we are planning to fully sustain the base level of operations via state support. Using these combined strategies, we envision SEFS and TESI core programs to become resilient and financially sustainable.

## Recommendations for Science Outreach Programs

A major recommendation for others looking to replicate a similar program to SEFS would be hiring coordinators, particularly former teachers, whose main responsibility is the facilitation of outreach programs. By having dedicated staff to coordinate and facilitate scientist classroom visits, it removes the burden on scientists and teachers. Since scientists do not need to organize the interaction, they can focus their time and energy on going through the SEFS training, planning their presentation, or practicing to make sure they are ready. Similarly, teachers are challenged to use any available time that they have most effectively. A Florida teacher communicated to the coordinators that before SEFS, between grading, lesson planning, connecting with parents, and more, it was too difficult to find the necessary time to arrange all aspects of a visit on their own. Although this teacher wanted to create these special experiences for students, they were unable to make them happen. As a result, if the resources allow, programs should hire or allocate a staff member to oversee the outreach process.

Another recommendation is to establish an organized database of scientists and teachers as well as a system to collect important metrics (such as grade level, topics, number of students reached, etc.). The latter also extends to implementing surveys for post-visit feedback and testimonials for promotional purposes that help the program grow. Further, an emphasis should be placed on relationship building between the scientist and teacher to create a mutually beneficial partnership. Both parties need to be comfortable with the process and not feel that participating is a daunting task. The program would therefore push out reminders, check-ins, and provide feedback to both the scientist and teacher. Both the teacher and scientist are also given best practices and ideas for engagement and successful interactions.

Additionally, all aspects of communications should be prioritized to make sure that the visit occurs effectively. This may include creating training for the scientists to cover research-based science communication tips and connecting with the teacher to discuss logistics and the teacher’s goal and vision for the visit (UF Thompson Earth Systems Institute, 2020; UF Thompson Earth Systems Institute, 2019). Although the SEFS program offers a training course on Canvas, this may be adapted or extended to other platforms as it makes sense for the specific target audience. This training aims to address scientist’s frequently asked questions, including specific topics to cover during the presentation, how often to communicate with their teacher counterpart, each party’s role, and more. By sharing clear information and communication from the beginning, these best practices will optimize the success of the partnerships and programs.

## How Scientists Partner with Us to Reach Florida K-12 Teachers and Students

There is an increasing emphasis on the relevance of what a scientist does and how they impact society. For example, the National Science Foundation requires “Broader Impacts” plans in grant proposals. Likewise, many scientific institutions expect their scientists to conduct outreach activities (MacFadden, 2019). Earth systems scientists interested in sharing their research to K-12 communities in Florida may collaborate with SEFS to reach their target audiences.

By partnering with SEFS for Broader Impacts, scientists gain access to a network of K-12 schools, teachers, and district leaders who have come to trust SEFS’s work. The team is composed of former K-12 educators who understand the unique and specific needs of classrooms and are well-versed in state learning standards. When a scientist partners with SEFS, the team will not only help match the scientist with the K-12 audience but will also help scientists craft their research expertise content in a way that is understandable, engaging, and fits into existing curricula.

## Concluding comments

Currently, in its second year of implementation, the SEFS program pilot has reached over 53,000 students during about 1,700 scientist classroom interactions involving over 850 teachers. The project holds promise as its efforts continue to grow with additional interested district leads, teachers, scientists, and as we continue to develop a sustainable funding model for the future. Our program tailors lessons and activities based on the individual needs of the teachers and their classrooms. The customization and direct collaboration between teacher and scientist are unique to the program and create a special experience for all parties involved.

Since the project is still in its first few years since inception, a limitation of the project is that there has not been a wide response in the evaluative efforts thus far. We realize this as a current shortcoming if we want to truly understand the impact that we are making on our teachers and their students. An IRB-approved STEP (Scientist-Teacher Educational Partnership survey) is in the beginning stages of implementation. The survey is designed to explore teachers’ and scientists’ perceptions of their partnership in designing and implementing instruction. It focuses on communication and planning, science teaching self-efficacy, and perceptions of student engagement and learning. Once additional responses have been submitted, an assessment will be able to be analyzed on the impact of the scientist-teacher relationships that are facilitated. Further, we are developing plans with research partners to measure impact on student academic growth from participating classrooms as a result of SEFS intervention efforts.

Overall we have learned many lessons and developed best practices over the first two years of SEFS. Over the next several years we will continue to refine programs to be grounded in student-centered learning and develop a sustainable funding model for the future. In doing so, we will be able to scale up and reach more schools, teachers, and students in Florida’s public schools, and ultimately be able to demonstrate impact. We envision that these activities will result in a better educated next generation of Floridians better poised to act responsibly in the 21st century, particularly as related to the future of the Earth and its natural systems.

## Acknowledgments

We thank all stakeholders for their participation in the Scientist in Every Florida School project. We are especially grateful for teachers who bring their enthusiasm and passion to our interactions. This work is part of the Scientist in Every Florida School Moonshot project funded by the UF Office of the President and Provost, Florida Museum, and Vice President for Research, with additional support provided by the Felburn and Smallwood foundations.

